# Multiple Targets, One Goal: Compounding life‐extending effects through Polypharmacology

**DOI:** 10.1101/2024.06.23.600269

**Authors:** K. Avchaciov, K. J. Clay, K. Denisov, O. Burmistrova, M. Petrascheck, P. Fedichev

## Abstract

Analysis of lifespan‐extending compounds suggested the most effective geroprotectors target multiple biogenic amine receptors. To test this hypothesis, we used graph neural networks to predict such polypharmacological compounds and evaluated them in *C. elegans*. Over 70% of the selected compounds extended lifespan, with effect sizes in the top 5% compared to the DrugAge database. This reveals that rationally designing polypharmacological compounds enables the design of geroprotectors with exceptional efficacy.

**Key takeaways:**

- The most effective known geroprotectors act by polypharmacological mechanisms.
- Graph neural networks predicted polypharmacological geroprotectors with a hit rate of 70%.
- The predicted polypharmacological geroprotectors are exceptionally effective.
- The predicted polypharmacological mechanism was experimentally confirmed.
- Rationally designing polypharmacological compounds results in geroprotectors with exceptional efficacy.

## Text

The genetic foundation of lifespan regulation is becoming increasingly well‐understood both within and across various species, revealing the key age‐associated genes. Small molecule drugs, the mainstay of the pharmaceutical industry, primarily modulate the activity of gene products—proteins, frequently targeting multiple pathways simultaneously by engaging multiple targets. Therefore, it is crucial to understand how the activity profiles of these molecules determine their life‐extending properties to optimize their anti‐aging effects.

The largest unbiased screen of biologically active molecules, particularly FDA‐approved drugs identified a significant cluster of drugs that extend lifespan by modulating neuroendocrine and neurotransmitter systems.^1,2^ We recognized that most G‐protein coupled receptors (GPCR) inhibitors interact with multiple structurally similar targets suggesting that polypharmacological action increases efficacy. To test this notion, we clustered target proteins based on sequence similarity (see methods). Consequently, known or predicted compounds target any protein within a cluster, were classified as “active” on that cluster (**Figure 1A**).

**Figure 1:**
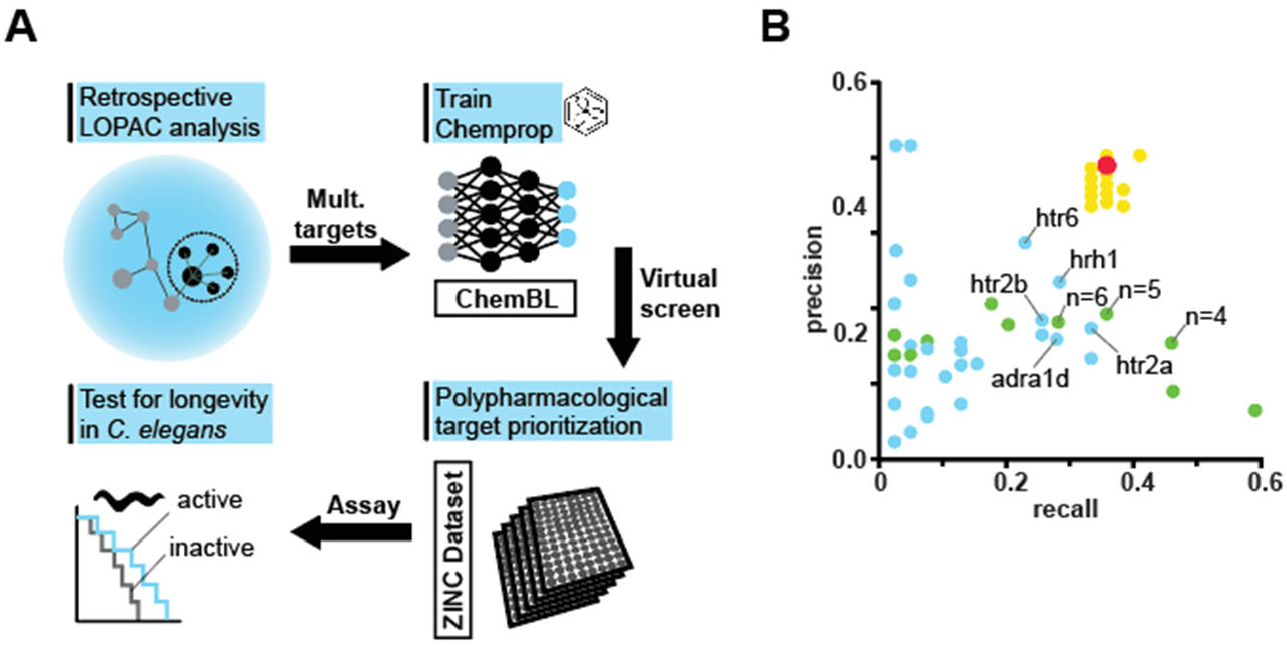
Computational modeling of lifespan extending molecules. **A)** Overview schematic of computational pipeline identifying polypharmacological compounds and validation in *C. elegans*. **B)** Precision‐recall scatter plot for the models implying either single cluster activity (blue dots), multiple clusters activity (green dots) and optimized combination of multiple clusters (yellow dots). Red indicates the model selected for the downstream analysis and experimental validation. Text labels indicate the leading target in the family of structurally related targets and the number of simultaneously identified activities for blue and green, respectively.

Next, we linked the lifespan extension property of a compound (LS+ label) to its activity against any single target by evaluating whether binding to any single target within a specified cluster significantly affects lifespan (**Table 1**). The best correlation (Matthews Correlation Coefficient [MCC] MCC=0.21) between activity and cluster, was the 5‐Hydroxytryptamine Receptor 6 (HTR6) gene. We characterized different aspects of model performance by standard precision and recall metrics. Precision characterized the model’s accuracy by measuring the proportion of correctly identified true lifespan‐extending drugs. Recall, or sensitivity, calculates the proportion of accurately predicted positive outcomes from all actual positives, evaluating the model’s ability to detect all true instances of lifespan extension.

**Table 1.**
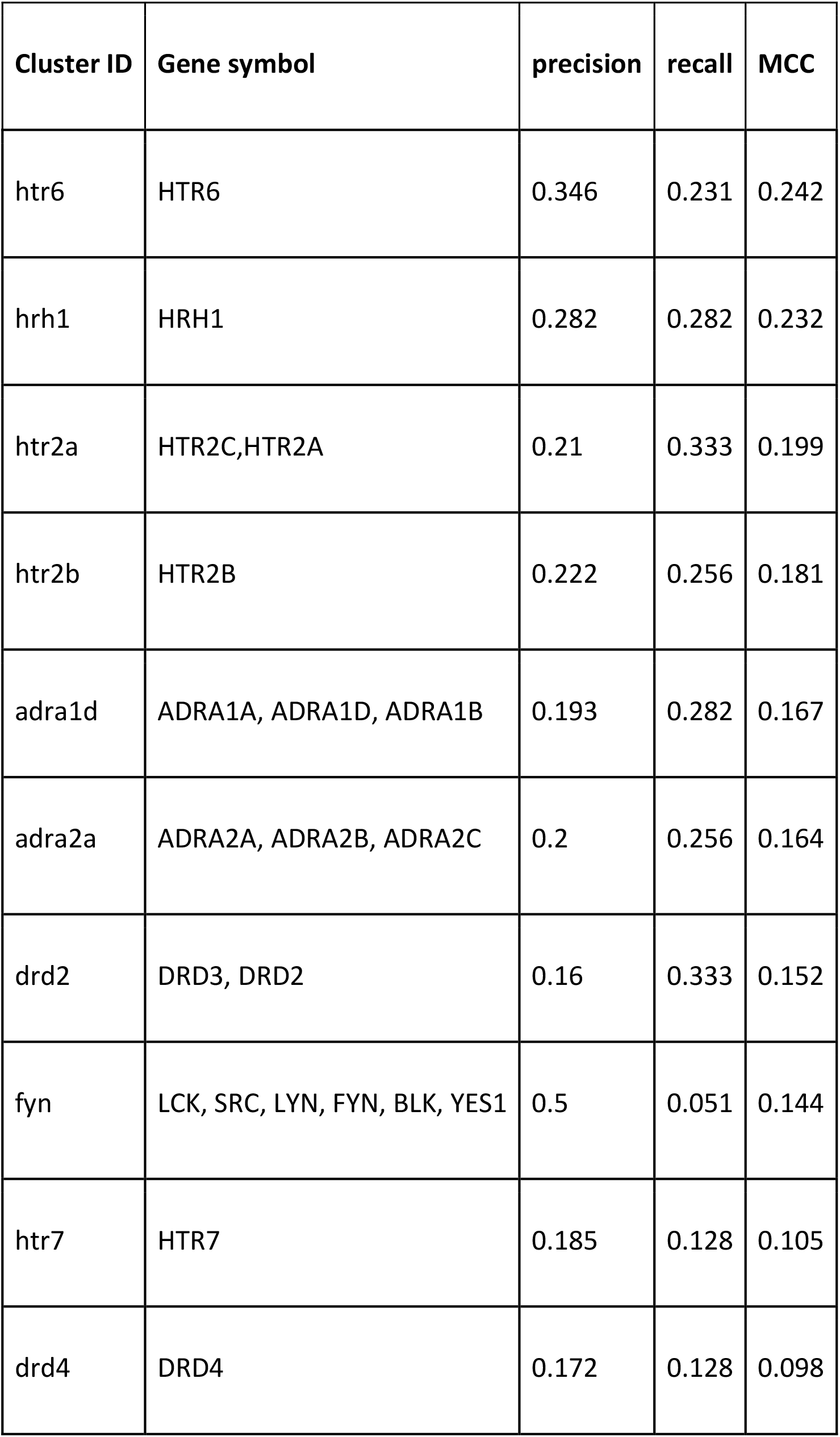
Top 10 predictions of extended lifespan label by an activity on a single cluster (MCC ‐ Matthews correlation coefficient)

We then investigated whether combining activities across different clusters could improve the prediction of the LS+ label. We developed a series of simple qualitative models that classify a molecule as LS+ based on its activity against combinations of up to five target classes. **Figure 1B** illustrates the recall‐precision relationships for models based on single cluster activity (blue dots) compared with models scoring molecules by the number of clusters with activity records (green dots). Finally, we characterized the models predicting the LS+ labels from a combination of activities against multiple target clusters (yellow dots).

The best performing model corresponded to a “polypharmacological” multi‐target model, necessitating activities across GPCRs clusters including DRD2, DRD3, HRH1, and HTR6 (**Figure 1B**, red dot). It exhibited significantly enhanced recall and precision compared to those based on activities against a single cluster (Fig 1B, precision 0.35, recall 0.23, MCC 0.24, see **Figure 1** and **Table 2** for the summary of all models analyzed).

**Table 2.** Prediction of extended lifespan label by an activity on a combination of clusters.

| Genes | rule | precision | recall | MCC |
| --- | --- | --- | --- | --- |
| [SHBG], [HTR6], [HRH1], [PGR,NR3C1,NR3C2,AR], [DRD3,DRD2] | 2 in<br>5 | 0.485 | 0.41 | 0.41 |
| [SHBG], [HRH1], [HTR1D,HTR1B], [PGR,NR3C1,NR3C2,AR], [DRD3,DRD2] | 2 in<br>5 | 0.483 | 0.359 | 0.382 |
| **[HTR6], [HRH1], [DRD3,DRD2]** | **2 in**<br>**3** | **0.467** | **0.359** | **0.373** |
| [SHBG], [HTR6], [HRH1], [NISCH], [DRD3,DRD2] | 2 in<br>5 | 0.467 | 0.359 | 0.373 |
| [SHBG], [HTR6], [HRH1], [DRD3,DRD2] | 2 in<br>4 | 0.467 | 0.359 | 0.373 |
| [HTR6], [HRH1], [LCK,SRC,LYN,FYN,BLK,YES1], [PGR,NR3C1,NR3C2,AR],<br>[DRD3,DRD2] | 2 in<br>5 | 0.467 | 0.359 | 0.373 |
| [HTR6], [HRH1], [NISCH], [PGR,NR3C1,NR3C2,AR], [DRD3,DRD2] | 2 in<br>5 | 0.467 | 0.359 | 0.373 |
| [HTR6], [HRH1], [NISCH], [LCK,SRC,LYN,FYN,BLK,YES1], [DRD3,DRD2] | 2 in<br>5 | 0.467 | 0.359 | 0.373 |
| [HTR6], [HRH1], [NISCH], [DRD3,DRD2] | 2 in<br>4 | 0.467 | 0.359 | 0.373 |
| [HTR6], [HRH1], [LCK,SRC,LYN,FYN,BLK,YES1], [DRD3,DRD2] | 2 in<br>4 | 0.467 | 0.359 | 0.373 |

To address the problem of sparse biological activity in the ChemBL database (most ligands have only been tested to bind a few targets), we leveraged modern machine learning algorithms to infer unmeasured activities for both known drugs and novel uncharacterized compounds. Specifically, we used Chemprop, a graph neural network utilizing the Directed Message Passing Neural Network (D‐MPNN) architecture,^3^ to analyze 29 targets and 50,000 activity records to predict optimized combinations of LS+ targets. We applied this model to a curated list of experimental and approved drugs to predict molecules active against at least two target–clusters simultaneously (Table 3).

**Table 3.**
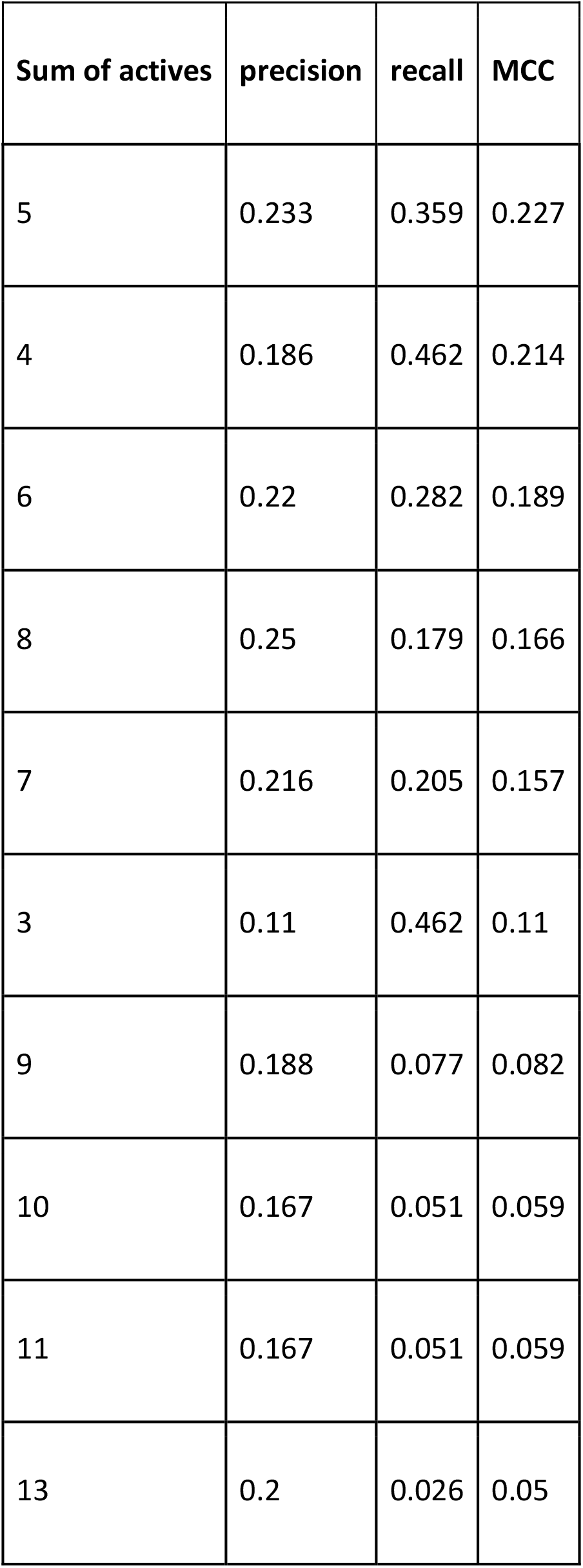
Prediction of extended lifespan label the by number of activity records per ligand.

In addition to that, we utilized the model to screen the ZINC library of purchasable compounds.^4^ Following medicinal chemistry filtering, we screened a set of 17 novel structures and 5 experimental or FDA‐approved drugs for lifespan extension in *C. elegans*.

Sixteen molecules significantly extended the lifespan of *C. elegans* at one or more doses (Supplementary **Table 1, Figure 2A, B**). The hit rate of over 70% (16/22) was approximately 1000‐fold higher than of an unbiased screen and more than sevenfold higher than the ∼9% hit rate expected from a set of compounds targeting biogenic amines (29/333 in the LOPAC screen^2^). Thus, a polypharmacology approach significantly outperforms selection based on pharmacology class. Similarly, the effect sizes of the current series of molecules were significantly larger than any previous screens. Of the 16 hits, 12 extended lifespan by over 40% and eight by more than 50%. The mean lifespan extension of the 16 hits exceeded 48%. The average lifespan extension annotated for all geroprotectors in DrugAge is ∼19%.^5^ Only 42 of the 973 geroprotectors annotated in DrugAge extend the lifespan of *C. elegans* by 44% or more, establishing our set in the top 5% of all known lifespan‐extending compounds.

**Figure 2:**
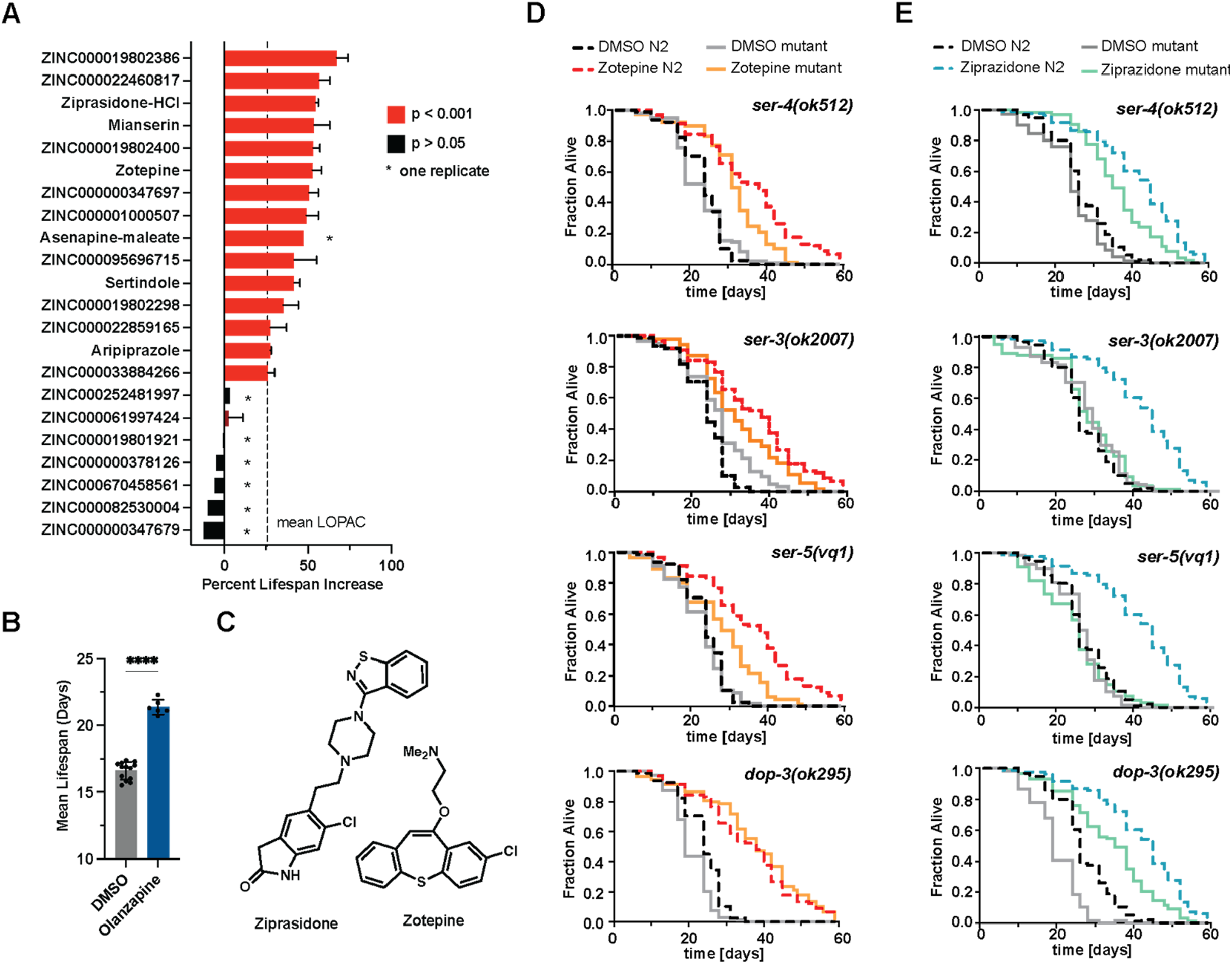
Computationally designed Geroprotectors target multiple G‐protein Coupled Receptors. **A)** Mean percent increase in lifespan extension observed at the optimal dose for two experiments for the 22 molecules designed. Mianserin was included as a blinded positive control. Error bars indicated mean ± SD. Mean lifespan extension observed for hits in the LOPAC screen is indicated by a dotted line to compare the effect sizes of between a phenotypic screening strategy and our current strategy. **B)** Olanzapine, used as an intra–plate positive control, consistently extended lifespan across the screen. Significance was determined by unpaired, two‐tailed, student’s t‐test where **** = p < 0.0001. Error bars indicated mean ± SD. **C)** Chemical structures of ziprasidone and zotepine. **D)** Graphs show survival plots of *C*.*elegans* wild type (N2), serotonin (*ser‐4, ser‐5*), octopamine (*ser‐3*) and dopamine (*dop‐3*) receptors treated with DMSO or Zotepine (10 μM). **E)** Graphs show survival plots of *C. elegans* wild type N2, serotonin (*ser‐4, ser‐ 5*), octopamine (*ser‐3*) and dopamine (*dop‐3*) receptors treated with DMSO or Ziprasidone (10 μM).

The maximum effect size observed in our study was induced by a compound identified by ML (+74%) for compound ZINC000019802386, which is structurally similar to Octoclothepin (Tanimoto similarity, T=0.45). There are no known activities against our GPCR targets for ZINC000019802386 but close analogs interact with DRD1, DRD2, DRD3, and HTR2A. This extension is comparable to the best‐performing compounds annotated in DrugAge.^5‐7^

To verify the polypharmacological properties of the hits, we selected two structurally distinct compounds, Ziprasidone and Zotepine, both active against targets in at least three of the four identified clusters (**Figure 2C**) and tested their effect in four mutant backgrounds most closely related to the annotated mammalian GPCRs: two serotonin receptor mutants, *ser‐4(ok512)* and *ser‐5(vq1)*, a dopamine receptor mutant *dop‐3(ok295)*, and an octopamine receptor mutant *ser‐3(ok2007)*. Octopamine is the invertebrate equivalent of noradrenaline, and SER‐3 is analogous to mammalian adrenergic receptors.^8‐10^ Ziprasidone and Zotepine were selected over the most active novel compound, ZINC000019802386, since the mammalian pharmacology of Ziprasidone and Zotepine has been established.^11^

As expected, the lifespan effects of both molecules depended on multiple GPCRs. Zotepine’s lifespan extension was contingent upon *ser‐3, ser‐4*, and *ser‐5* (**Figure 2D**). Although Zotepine still extended the lifespan in the mutants *ser‐3(ok2007), ser‐4(ok512)*, and *ser‐5(vq1)*, the effect was less pronounced compared to the wild‐type N2, suggesting that binding to each receptor contributed partially and additively to the overall effect. However, the dopamine receptor *dop‐3* was not necessary for the longevity induced by Zotepine. Similarly, Ziprasidone’s effect on extending the lifespan of *C. elegans* also required the same three GPCRs: *ser‐3, ser‐4*, and *ser‐5* (**Figure 2E**). Like Zotepine, Ziprasidone exhibited a reduced longevity effect in the *ser‐4(ok512)* mutant. Interestingly, unlike Zotepine, the lifespan extension by Ziprasidone was eliminated in the *ser‐3(ok2007)* and *ser‐5(vq1)* mutants, indicating a critical dependency on these receptors.

A simple model in which GPCRs act as binary “on‐off” switches, with ligands blocking signaling by the different receptors, only explains the interaction of Zotepine with the receptor mutants. In this binary model, blocking signaling by each receptor modulates lifespan independently, partially contributing to the total effect size. This model does not explain the result for Ziprasidone. More recent models suggest modulation of GPCRs by “ligand‐bias”, in which binding of the ligand causes the GPCR to recruit distinct downstream effectors such as G‐proteins, G‐proteins receptor kinases, or arrestins favoring one signaling modality over another adding further complexity to the interpretation of polypharmacological effects.^12,13^

Thus, a polypharmacological ligand may have different functional selectivity or ligand bias for each individual receptor.^14^ While the affinity of a ligand recruits it to a receptor, ligand‐bias modulates the downstream signaling resulting in outcomes for ligands binding to the same receptor. Zoptepine and Ziprasidone require the same receptors for their full lifespan extension but the receptors’ quantitative contributions to the overall longevity effect differ. This observation suggests that the two compounds differ in their downstream effector response. Similar considerations also explain why GPCR ligands often do not phenocopy receptor knockout phenotypes and *vice versa*.

In this study, we deviated from target–based drug discovery and explored the potential of a polypharmacological strategy to extend lifespan by generating small molecules that target multiple targets simultaneously. Instead, we followed our observation that many of the most effective lifespan‐extending compounds demonstrated polypharmacological activity profiles.^8,15,16^ To overcome the complexity of rational design of polypharmacological compounds, we employed a combination of statistical and machine learning tools to leverage insights from a large phenotypic lifespan screen. The resulting computational strategy identified polypharmacological compounds whose validation confirmed the power of polypharmacology in the design of geroprotectors by: (I) an exceptional hit rate of over ∼70%, (II) an exception effect size in extending lifespan by over 40% compounds across the entire set and (III) multi‐target effects of the selected compounds.

The proposed approach markedly differs from the prevalent method in current drug discovery, which identifies molecules with high selectivity against a single therapeutic target. Until recently, polypharmacological action was typically accidental and often considered undesirable for two reasons. First, the lead‐to‐drug optimization becomes far more complex when multiple targets are involve.^17^ Second, targets are often evaluated based on their genetics, mostly single gene knockouts or alleles, with far less data available on how different alleles interact. Consequently, polypharmacological design principles are challenging to implement in conventional drug discovery pipelines, in part, because “off‐target effects” have often been found detrimental. However, retrospective analyses of successful drugs show that their therapeutic efficacy or safety often depends on multiple targets, highlighting that unintended polypharmacology contributes to the success of a drug.^18‐20^ Our study establishes that computational strategies can solve the increased complexity of polypharmacological strategies and validate their complex predictions in vivo.

The only shortcoming in the current study was that the most novel structures did not extend lifespan. This observation could be explained in two ways. First, the efficacy of a compound in an experiment may depend on more than mere target engagement but also require suitable absorption, distribution, metabolism, excretion, and toxicity (ADMET) properties. None of the attributes were used in our computational pipeline and hence the performance of our models may suffer when we deviate from well characterized structural motifs shared with approved drugs.

Second, large neural networks, such as Chemprop used here, operate as “black boxes” that need to rely on limited experimental data typically available in biomed datasets. Our demonstration provides evidence that more often than not such systems operate as search engines in the chemical space and tend to favor compounds structurally similar to those active in the training data. The current limitations may be overcome as more data becomes available for training models, network architectures improve, and more aggressive train/test set splits could be employed to achieve better generalization.

In summary, we produced evidence suggesting that selecting geroprotectors based on their polypharmacological activities against targets implicated in regulation of longevity process may be useful in identifying compounds with significant effects on lifespan. We demonstrate that developing polypharmacological geroprotectors can be assisted by modern ML tools and may help enhance efficacy in lifespan extension. While the neural network in this study did not identify novel pharmacophores, this outcome is secondary within the context of this work. We are optimistic that more advanced ML techniques will eventually reach the generalization capabilities sufficient for the systematic identification of new chemical entities that utilize polypharmacology for greater geroprotective effects.

## Acknowledgements

PF and AK thank Dr. Andrey Tarkhov and Simon Steshin from Gero (at the time of writing) for discussions, suggestions and technical assistance. Part of this work was funded by R01 AG067331 and R01 AG069206 (to M.P).

AK, KD, OB, and PF are employed by Gero and declare no conflict of interest.

## Methods

### Statistical analysis of LOPAC library screening

The LOPAC dataset was built from the Supplementary materials from Ye et al.^1^ The compound SMILES were retrieved from Pubchem using PUG‐REST.^21^ From the SMILE string we obtained all InchiKeys, including the InchiKey for the compound itself and the InchiKeys for the parent compounds. The InchiKey was used to get activity data from the ChEMBL web services.^22^ In case no activity records were found for the precise InchiKey match, we search records by the first 14 characters of the InchiKey string. We obtain activities with the threshold in the minimal pChEMBL value equals 5 for 1029 of 1275 total compounds from the LOPAC dataset.^23^ Next we selected binding (B) and functional (F) assays with an inhibition type outcome (Ki, IC50, Kd, etc). ChEMBL targets were clustered by protein sequence similarity using MMseqs2 with the sequence identity threshold of 0.5.^24^ The activity values were aggregated within all targets of one cluster with the min function and then binarized at the threshold of pChEMBL=7 (corresponding to 100 nM in standard units). Hereinafter, we defined target as a cluster of similar proteins (see Supplementary **Table 1**) and activity on this target defined by the minimal value of pChEMBL from proteins in it. This results in 1256 LOPAC compounds with known structure and 281 activity targets, among which only 594 compounds have at least one active target.

The binary activity label is used for predicting a lifespan extension (LS+) label. We used standard metrics: precision, recall, and the Matthews Correlation Coefficient (MCC), each of which characterizes different aspects of model performance.^25^ For instance, precision measures the proportion of accurately predicted positive outcomes from all predicted positives, assessing the model’s accuracy in identifying true lifespan‐extending drugs. Recall, or sensitivity, calculates the proportion of accurately predicted positive outcomes from all actual positives, evaluating the model’s ability to detect all true instances of lifespan extension. The MCC offers a comprehensive evaluation of binary classification by accounting for both positive and negative outcomes, making it especially useful in imbalanced datasets. It ranges from ‐1 (inverse prediction) to +1 (perfect prediction).

For the models based on the number of activity records per ligand, we calculate the number of activities and produce a binary label if this number is greater or equal to N, where the number N varies from 2 to 20.

For the model based on optimal selection of combinations of targets, we generated combinations of targets (from 2‐length to 5‐length tuples) and calculated the binary labels at different thresholds of simultaneously active targets varying from 1 to length of the tuples. It can be expressed as a logic rule “***n*** in ***M***”, where ***n*** is the number of simultaneously active targets, and ***M*** is the size of a tuple. For example, the best model in this work is “**2 in 3”** for the 3‐length tuple of targets ([DRD2,DRD3], [HRH1], [HTR6]), which requires at least two targets to be non‐zero (**Supplementary Table 2**).

### Graph neural network for predicting molecular interactions

We employed the graph neural network Chemprop to predict the binding affinities of small molecules from the CHEMBL database.^3,22^ Chemprop, a Directed Message Passing Neural Network, learns molecular properties from the graph structure of molecules, where atoms serve as nodes and bonds as edges. For each molecule, we reconstructed its molecular graph from its SMILES string and identified the atoms and bonds using the RDKit open‐source package.^26^ We then initialized a feature vector for each molecule based on computable features listed in the table [**Supplementary_table_[rdkit_features]**]. List of descriptors was modified from the original Chemprop paper by removing highly correlated features for speed improvement.

The model performs a series of message passing steps, aggregating information from neighboring atoms and bonds to understand the local chemistry. This information, combined with precomputed molecular features, is processed through a feed‐forward neural network, which predicts the property of interest. In our case, we solved the multiclass classification problem of whether a molecule binds to one of target GPCR proteins listed in the **Supplementary_table_[chemprop_targets]**. We used the same method of retrieving binary labels for each target as described in “Statistical analysis of LOPAC library screening”. The unknown activity values were excluded from the loss evaluation during training.

We enhanced the original Chemprop implementation by integrating DenseNet‐like skip connections into the feedforward segment of the network. In this configuration, each layer sequentially receives the concatenated activations from all preceding layers, enabling every neuron to access the learned features comprehensively.^27^ Train‐validation‐test split was performed with Butina algorithm^28^ and the cutoff 0.8 for Tanimoto similarity computed over RDKit Fingerprints and in the ratio 0.8/0.1/0.1. We performed a simple hyperparameter search. Optimal parameters were selected using grid search technique using the ROC‐AUC metric function on the test set. This procedure led us to the selection of an architecture described in the **Supplementary_table_[chemprop_params]**. Metrics on the test set at the end of the training procedure are listed in the **Supplementary_table_[chemprop_metrics]**.

We scored the ZINC in‐stock subset to predict the activities of the molecules against the selected GPCR targets and chose molecules with required predicted polypharmacological effect. We further applied QED\ref[] filter and obtained 66 candidates for the experiment see **[Supplementary_table_ZINC**]. To reduce the experimental work, we used Butina clustering algorithm to select structurally dissimilar compounds. At the end, we reduced the list to 16 potentially active and non‐redundant compounds.

### *C. elegans* strains

The Bristol strain (N2) was used at the wild‐type strain. The following strains used in this study were obtained from the Caenorhabditis Genetic Center (CGC; Minneapolis, MN): RB1631 [s*er‐3(ok2007)]*, AQ866 *[ser‐ 4(ok512)]*, and BZ973 *[dop‐3(ok295)]*. The strain VV212 [*ser‐5(vq1)]* was generated in Perez‐Gomez et. al.^10^

### Lifespan assays

Lifespan assays were conducted in liquid culture 96 well plates as outlined in Clay et. al.^29^ Each permutation (strain‐drug) was tested in trials of 50‐100 animals in each trial. These numbers have sufficient power to detect differences in lifespan of 20% or more in more than 95% of the time.^30^ Animals were treated with drugs on day 1 of adulthood to avoid any developmental effects. Lifespan experiments were conducted blind. Statistical analysis was performed using the Mantel‐Haenzel version of the log‐rank test and data are summarized in **Supplementary_table_[Lifespan]**.

### Supplementary Tables

Supplementary_table_[lifespan]: Summary of lifespan statistics

Supplementary_table_[drug_candidates]: Selected candidates for screening

Supplementary_table_[chemprop_targets]: Molecular targets with gene symbols

Supplementary_table_[rdkit_features]: Molecular features and their RDKit implementation names

Supplementary_table_[chemprop_params]: Model hyperparameters and training parameters

Supplementary_table_[chemprop_metrics]: Classification metrics on test data subset

Supplementary_table_[ZINC]: List of selected compounds and ZINC parameters

